# Evaluation of the α-synuclein PET radiotracer (d_3_)-[^11^C]MODAG-001 in pigs

**DOI:** 10.1101/2022.02.05.479231

**Authors:** Nakul Ravi Raval, Clara Aabye Madsen, Vladimir Shalgunov, Arafat Nasser, Umberto Maria Battisti, Emily Eufaula Beaman, Morten Juhl, Louise Møller Jørgensen, Matthias Manfred Herth, Hanne Demant Hansen, Pontus Plavén-Sigray, Gitte Moos Knudsen

## Abstract

**Background:** A positron emission tomography (PET) radiotracer to neuroimage α-synuclein aggregates would be a crucial addition for early diagnosis and treatment development in disorders such as Parkinson’s disease, where elevated aggregate levels is a histopathological hallmark. The radiotracer (d_3_)-[^11^C]MODAG-001 has recently shown promise for visualization of α-synuclein pre-formed fibrils (α-PFF) in rodents. We here test the radiotracer in a pig model where proteins are intracerebrally injected immediately before scanning. Four pigs were injected in one hemisphere with 150 µg α-PFF, and in the other hemisphere, either 75 µg α-PFF or human brain homogenate from either dementia with Lewy bodies (DLB) or Alzheimer’s disease (AD) was injected. All pigs underwent one or two (d_3_)-[^11^C]MODAG-001 PET scans, quantified with the non-invasive Logan graphical analysis using the occipital cortex as a reference region.

**Results:** The α-PFF and AD homogenate injected brain regions had high uptake of (d_3_)-[^11^C]MODAG-001 compared to the occipital cortex or cerebellum. BP_ND_ values in 150 µg α-PFF injected regions was 0.78, and in the AD homogenate injected regions was 0.73. By contrast, the DLB homogenate injected region did not differ in uptake and clearance compared to the reference regions. The time-activity curves and BP_ND_ values in the 150 µg and 75 µg injected region of α-PFFs show a dose-dependent effect, and the PET signal could be blocked by pretreatment with unlabeled MODAG-001.

**Conclusion:** We find that both α-PFF and AD brain homogenates give rise to increased binding of (d_3_)-[^11^C]MODAG-001 when injected into the pig brain. Despite its limited specificity for cerebral α-synuclein pathology, (d_3_)-[^11^C]MODAG-001 shows promise as a lead tracer for future radiotracer development.

## Background

Parkinson’s disease (PD), dementia with Lewy bodies (DLB), and multiple system atrophy (MSA) are histopathologically characterized by progressive nigrostriatal, limbic and neocortical neurodegeneration and aggregation of the intracellular presynaptic protein α-synuclein [1–3]. These diseases are collectively known as α-synucleinopathies [4]. Patients with PD or DLB have α-synuclein-rich neuronal inclusions called Lewy bodies and Lewy neurites, predominantly in the substantia nigra in PD and throughout the cerebral cortex in DLB [5]. On the other hand, patients with MSA show filamentous aggregates in oligodendrocytes and neurons [6]. However, it is yet unknown to which extent α-synuclein aggregates contribute to neurodegeneration (Wong and Krainc 2017), and further, the clinical diagnosis of a PD or PD+ disorder is difficult, particularly in the early phases [7]. In drug-naive patients with subtle clinical parkinsonian motor symptoms, dopamine transporter neuroimaging has high sensitivity and specificity in distinguishing between patients with and without striatal neurodegeneration [8] but access to a neuroimaging tool to specifically assess α-synuclein aggregates would be a highly valuable addition.

Positron emission tomography (PET) has proven valuable for the detection of amyloid-β and tau protein aggregates and is used for differential diagnosis and drug development evaluation for neurodegenerative conditions such as Alzheimer’s disease (AD) [9]. PET imaging of α-synuclein would be advantageous for, e.g., early disease detection, differential diagnosis, and monitoring disease progression of synucleinopathies. In addition, the field is moving towards early eradication of α-synuclein aggregates as a promising therapeutic strategy in α-synucleinopathies, an approach that would require in vivo imaging for clinical application. As of today, no clinically validated PET radioligand exists for imaging α-synuclein [10, 11].

Several attempts to develop a suitable radioligand for α-synuclein have been made, and some tracers looked promising in rodents [12–16]. One of these is the diphenylpyrazole derivative [^11^C]MODAG-001/(d_3_)-[^11^C]MODAG-001 [12]. It was developed from the lead structure anle138b, a compound with therapeutic properties in PD and MSA rodent models due to its binding characteristics to α-synuclein aggregates [17, 18]. Anle138b and its derivatives, like [^3^H]/[^11^C]MODAG-001, have undergone extensive in vitro and rodent biodistribution experiments [12, 19]. (d_3_)-[^11^C]MODAG-001 showed the most promise as a candidate radioligand for detecting α-synuclein aggregates due to its high affinity, good brain penetration, and ability to detect α-synuclein pre-formed fibril (α-PFF) in a protein deposition rat model [12].

In the present study, we test and characterize (d_3_)-[^11^C]MODAG-001 in a large animal model. To evaluate binding characteristics to α-synuclein aggregates, we use a pig model where α-PFF is intracerebrally injected (ICI) immediately before scanning, creating an artificial target brain region [20]. We assess the sensitivity of the radioligand to identify α-synucleinopathy DLB human brain homogenate injected in the pig brain. The binding selectivity to α-synuclein is assessed by comparison to a brain region where amyloid and tau pathology-rich AD human brain homogenate is injected.

## Methods

### Radiochemistry

Precursor and reference compound for (d_3_)-[^11^C]MODAG-001 were prepared as previously described [12]; see supplementary information for more details. (d_3_)-[^11^C]MODAG-001 was obtained by reductive amination of desmethyl precursor with [^11^C]CH_2_O. The radioactivity yield was 650±297 MBq (mean±SD) (n=9, range 250-1214) after 70 min of synthesis time. The radiochemical purity of the formulated tracer was >95%. Radiochemical conversion from trapped [11C]CH3I was 45±2 % (n=6). Molar activity at the end of the radioligand synthesis was on average 28.14±5.3 GBq/μmol.

### Animals

We included four female domestic pigs (crossbreed of Landrace, Yorkshire, and Duroc) weighing 27±1 kg and aged 10-11 weeks (Table 1). Before any experiments, pigs were sourced from a local farm and acclimatized for 7-10 days in an enriched environment.

**Table 1.**
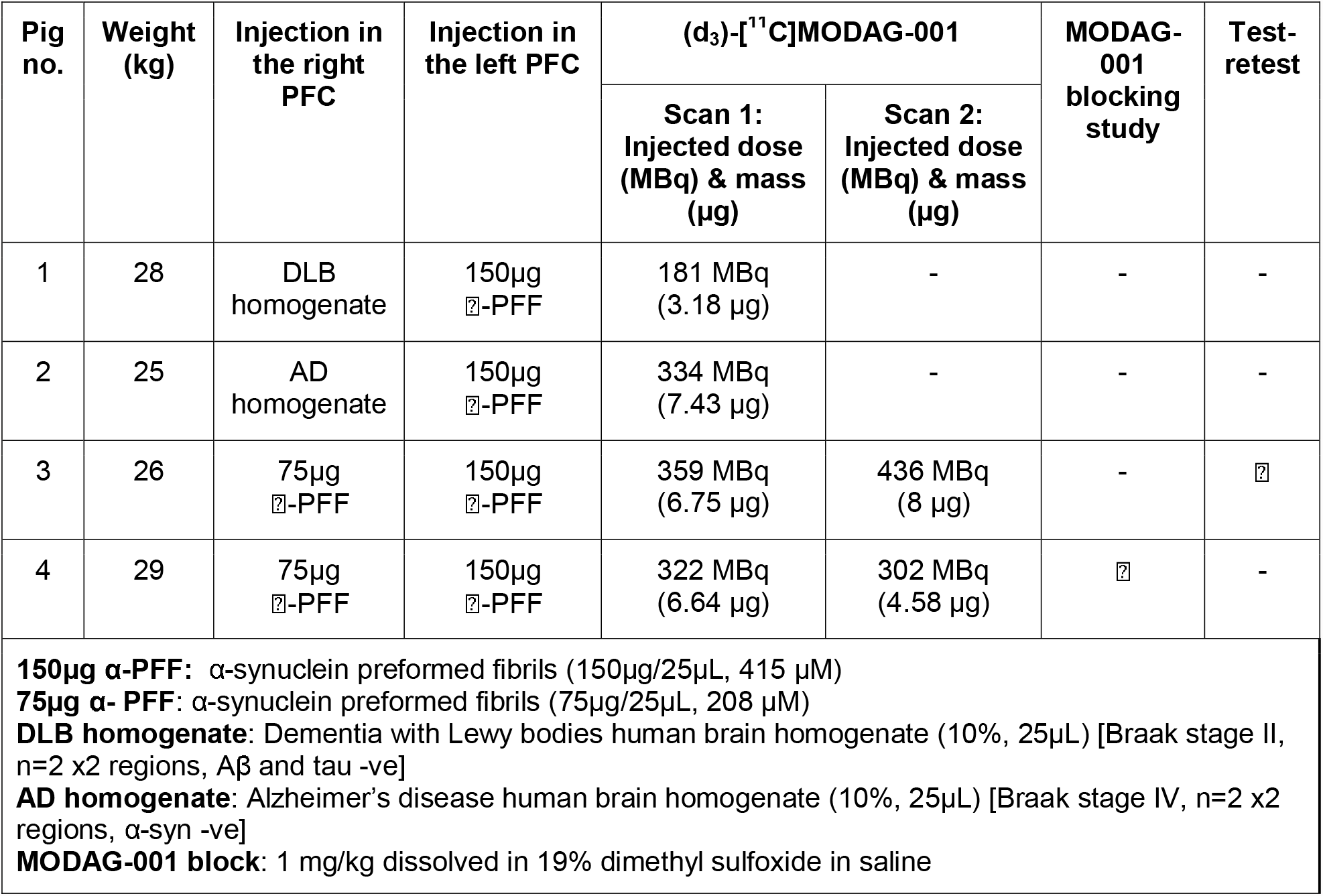
Pig characteristics: Body weight, injectate in the PFC, injected dose/mass of (d_3_)-[^11^C]MODAG-001, and availability of blocking and test-retest scans.

### Preparation and surgical procedure

A detailed description of the preparation, anesthesia, surgery, and transport has previously been described [21, 22]. Briefly, anesthesia was induced with an intramuscular (IM) injection of Zoletil mixture and maintained with 10-15 mg/kg/h propofol intravenous (IV) infusion. Analgesia was achieved with 5 µg/kg/h fentanyl IV infusion. Endotracheal intubation allowed for ventilation with 34% oxygen in normal air at 10-12 mL/kg. The left and right superficial mammary veins, ear veins, and femoral arteries were catheterized for venous and arterial access. The animals’ heart rate, blood pressure, peripheral oxygen saturation (SpO_2_), end-tidal CO_2_ (EtCO_2_), blood glucose, and temperature were monitored throughout the scan. Using a modified stereotactic approach [20], pigs were intracerebrally injected into the medial prefrontal cortex (mPFC) with 25 µL of 3 or 6 mg/mL α-PFF (molecular weight of monomer: 14,460 Da, corresponding to 208 µM or 415 µM) (produced at H. Lundbeck A/S, Copenhagen, Denmark), AD human brain homogenate (10% homogenate in saline [α-synuclein aggregates -ve]) or DLB human brain homogenate (10% homogenate in saline [amyloid-β and tau aggregates -ve]), as outlined for each pig in Table 1. The post-mortem human brain homogenates used in this study are the same as described previously [20], namely a homogenate mixture of two regions (frontal and temporal) from 2 different patients with each disease (i.e., AD and DLB). In a previous study, the injection target point in the mPFC: 8, 25, 14 mm in X, Y, Z coordinates relative to bregma was validated [20]. After the surgical procedure, the animals were transported to the scanner facilities.

### PET scanning protocol

Pigs were PET-scanned either once or twice (same day) in a Siemens high-resolution research tomograph (HRRT) scanner (CTI /Siemens, Malvern, PA, USA). (d_3_)-[^11^C]MODAG-001 was injected as a rapid bolus (∼20 seconds) through one of the superficial mammary veins (IV), and PET data were acquired over 121 min. Molar activity at the time of injection was 19.0±2.1 GBq/μmol (injected dose and mass in Table 1). Pig 3 received a test-retest on the same day. In Pig 4, we perform a self-blocking study with 1 mg/kg non-deuterated unlabelled MODAG-001. Unlabelled MODAG-001 (29.1 mg) was dissolved in 40 mL of saline with 19% dimethyl sulfoxide to ensure full solubility and injected IV over 15 min starting ∼6 min before the injection of (d_3_)-[^11^C]MODAG-001.

### Blood sampling and radio-HPLC analysis

Radio-HPLC analysis of plasma samples were performed in Pig 4 for both baseline and block scans. Manual arterial blood samples were drawn at 1.5, 5, 20, 40, and 60 min after injection. Samples were also drawn at 90 and 120 min, but data is not shown due to low and noisy radioactivity counts. Pig 4 also received a third injection of (d_3_)-[^11^C]MODAG-001 (180 MBq, 3.74 μg) to assess radiometabolites crossing the blood-brain barrier for which a blood and brain sample was acquired at 15 min and 22 min post tracer injection. A blood sample was drawn before injection of 20 mL pentobarbital/lidocaine for euthanasia. Immediately after, the skull was exposed, and the occipital bone was sawed open. A small brain sample from the occipital cortex was excised and rinsed in saline to remove excessive blood. Radiolabeled parent and metabolite fractions were determined in plasma and brain tissue using isocratic elution, as previously described [12], but with some modifications (details in Supplementary information).

### *In vitro* methodologies

After the last scanning, the animals were euthanized by IV injection of 20 mL pentobarbital and lidocaine. After euthanasia, the brains were removed, snap-frozen with powdered dry-ice, and stored at −20 °C until further use. Intracerebrally injections were confirmed using fluorescence immunohistochemistry; procedure and results are available in the supplementary data.

### PET data reconstruction and preprocessing

PET scans were reconstructed using ordinary Poisson 3D ordered subsets expectation-maximization, including modeling the point-spread function, using 16 subsets, ten iterations, and standard corrections [23]. Attenuation correction was performed using the MAP-TR μ-map [24]. Emission data were binned into time frames of increasing lengths: 6 × 10 s, 6 × 20 s, 4 × 30 s, 9 × 60 s, 8 × 120 s, 4 × 180 s, 2 × 240 s, 1 × 300 s, 1 × 360 s, 1 × 420 s, 1 × 600 s, 1 × 900 s, and 1 × 1680 s. Each frame consisted of 207 planes of 256 × 256 voxels of 1.22 × 1.22 × 1.22 mm in size. Brain parcellation was performed according to our previously published automatic PET-MR pig brain atlas method [25]. The input for the methodology was frame-length weighted, summed PET images of the total scan time (0–120 min). Time-activity curves (TACs) from the neocortex, occipital cortex, temporal cortex, cerebellum (here defined as without vermis), and injection regions were extracted for the present study. The regions of the injection sites were delineated as described in our previous study [20], while all other regions were part of the Saikali atlas [26] modified for PET [25].

### Pharmacokinetic modeling

For image quantification, we used the non-invasive Logan graphical analysis [27] with the occipital cortex and cerebellum as reference regions. In order to estimate the average k_2_ over R_1_ ratio (k_2_’), we applied the simplified reference tissue model (SRTM) [28] to high binding regions (i.e., α-PFF injected regions) and calculated k_2_’. For the non-invasive Logan plot, we chose the threshold time, t*, of 23 min (last 15 frames) since it showed the lowest average maximum percentage of variance. BP_ND_ values estimated using the occipital cortex as a reference region were more stable than those derived using the cerebellum. These are therefore presented in the results section below.

All kinetic modeling was performed using the *“kinfitr” package* (v. 0.6.1) (Matheson, 2019; Tjerkaski et al., 2020) in R (v. 4.0.2; “Taking Off Again,” R core team, Vienna, Austria). For the pig that received a test-retest scan, we calculated the % test-rest change using Equation 1. For the pig that received a baseline-block scan, we calculated the % blocking in the α-PFF injected regions using Equation 2.

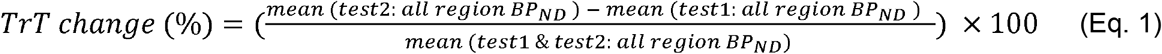

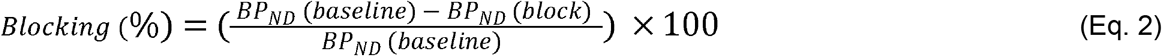

Regional radioactivity concentration (kBq/mL) was normalized to injected dose (MBq) and corrected for the animal weight (kg) to provide standardized uptake values (SUV, g/mL) used in graphical plots in Figures 1 and 3. PMOD 3.7 (PMOD Technologies, Zürich, Switzerland) was used to visualize and create all representative PET images (Figure 1 and 3), which are summed images over the entire period of the scan (0-121 min) with the “Triangle” PMOD pixel interpolation function; for more details see “https://www.pmod.com/files/download/v31/doc/pbas/4145.htm”. Graph-Pad Prism (v. 9.2.0; GraphPad Software, San Diego, CA, USA) was used for data visualization.

**Figure 1.**
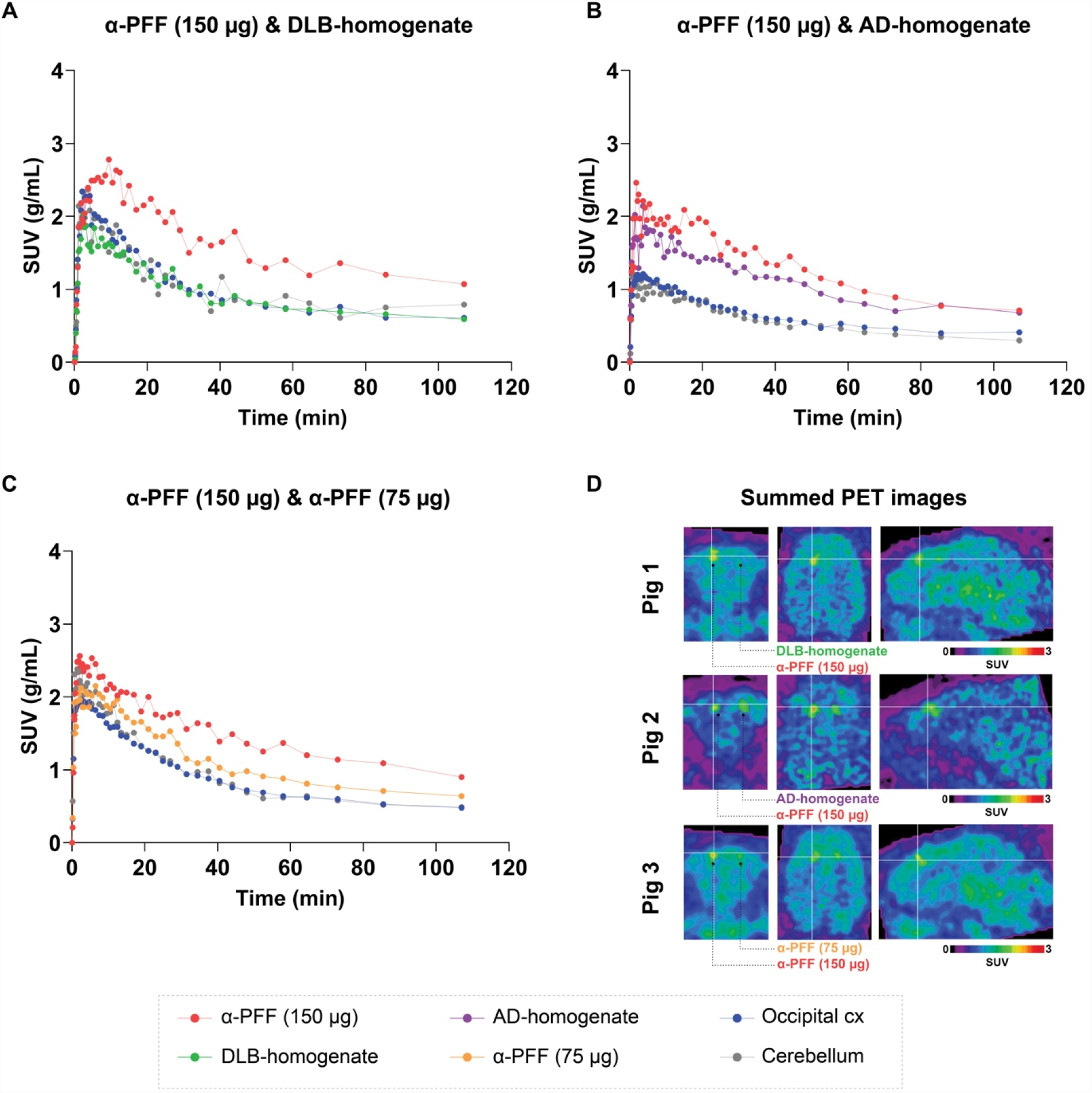
Regional TACs of (d_3_)-[^11^C]MODAG-001 in pigs injected with 150 µg α-PFF and A) DLB homogenate, B) AD homogenate and C) 75 µg α-PFF. TACs for the two reference regions, ie, the occipital cortex and cerebellum, are also shown. D) SUV-scaled PET images from representative TACs.

## Results

### Brain uptake and kinetics of (d_3_)-[^11^C]MODAG-001

We observed high brain uptake (∼ 2.5 SUV) and a relatively quick radioligand wash-out after (d_3_)-[^11^C]MODAG-001 injection. The plasma kinetics of (d_3_)-[^11^C]MODAG-001 were relatively fast, with approximately 10% of the parent radioligand remaining in plasma after 20 min (Supplementary Figure 1). Regions with either 150 µg (n = 4) or 75 µg (n = 2) α-PFF and AD homogenate (n = 1) had higher radioactivity retention (Figure 1A-C and Figure 2A) compared to the occipital cortex and cerebellum. By contrast, the DLB homogenate region (n = 1) TAC behaved essentially as background tissue radiotracer retention (Figure 1A). Almost identical TACs were seen in the pig with test-retest scans (Supplementary Figure 2). In a pig euthanized 15 min after tracer injection, 10.8% of (d_3_)-[^11^C]MODAG-001 parent compound remained in the plasma while 56.1 % parent compound remained in brain homogenate from the occipital cortex (Supplementary Table 1). The remaining signal from the plasma and brain came from polar and non-polar radiometabolites (Supplementary Table 1).

**Figure 2.**
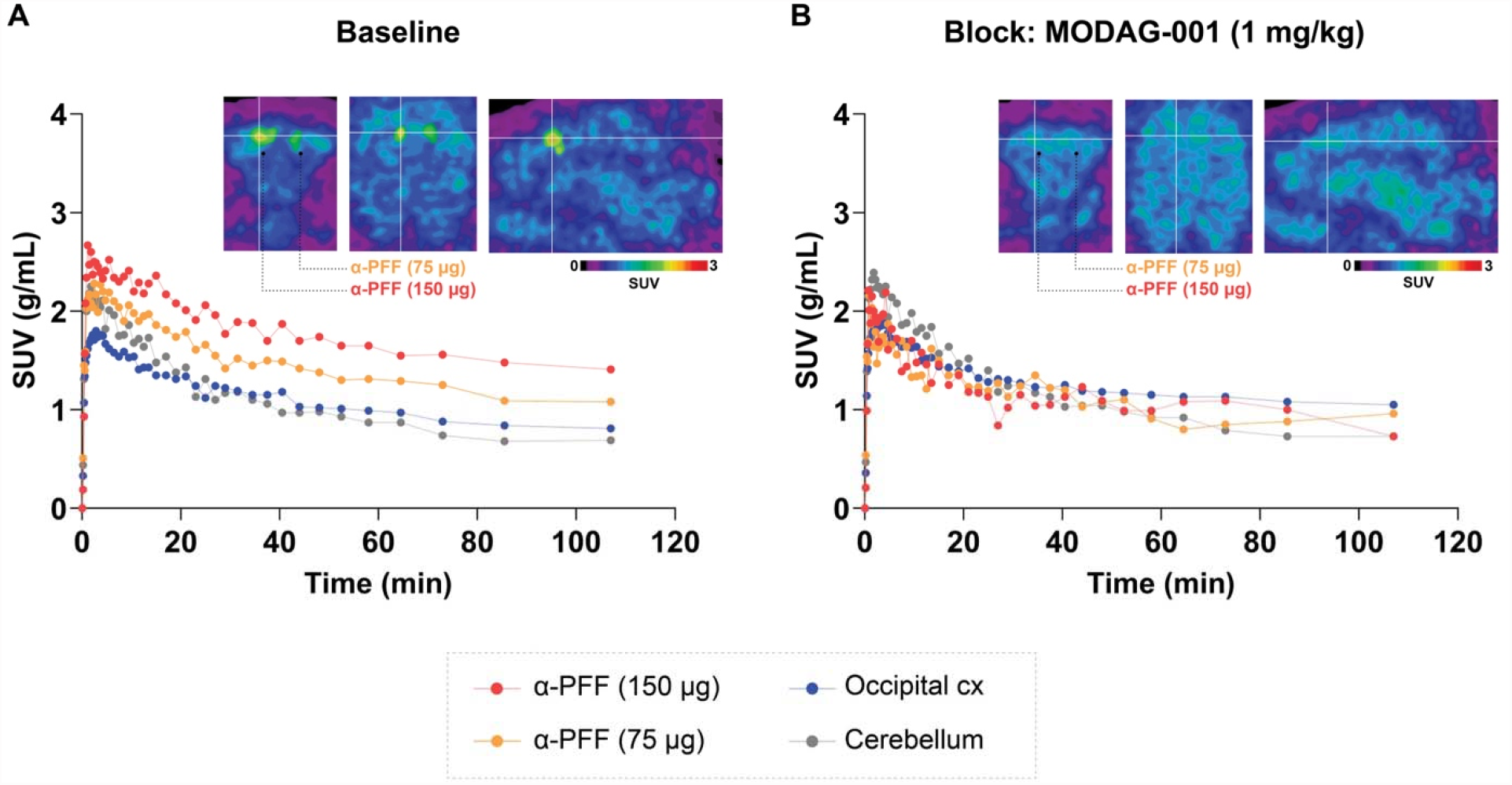
(d_3_)-[^11^C]MODAG-001 baseline and block. TACs and SUV scaled PET images A) (d_3_)-[^11^C]MODAG-001 baseline and B) (d_3_)-[^11^C]MODAG-001+ MODAG-001 (1 mg/kg) block scan from a pig with 150 µg and 75 µg α-PFF.

### Blocking experiment using MODAG-001

Pretreatment with 1 mg/kg MODAG-001 shortly before the injection of (d_3_)-[^11^C]MODAG-001 significantly reduced the radioactive signal in the 150 µg and 75 µg α-PFF regions, which showed substantially faster radioligand kinetics than the regional baseline TACs (Figure 2B); the TACs became comparable to those in the occipital cortex and cerebellum (Figure 3B).

**Figure 3.**
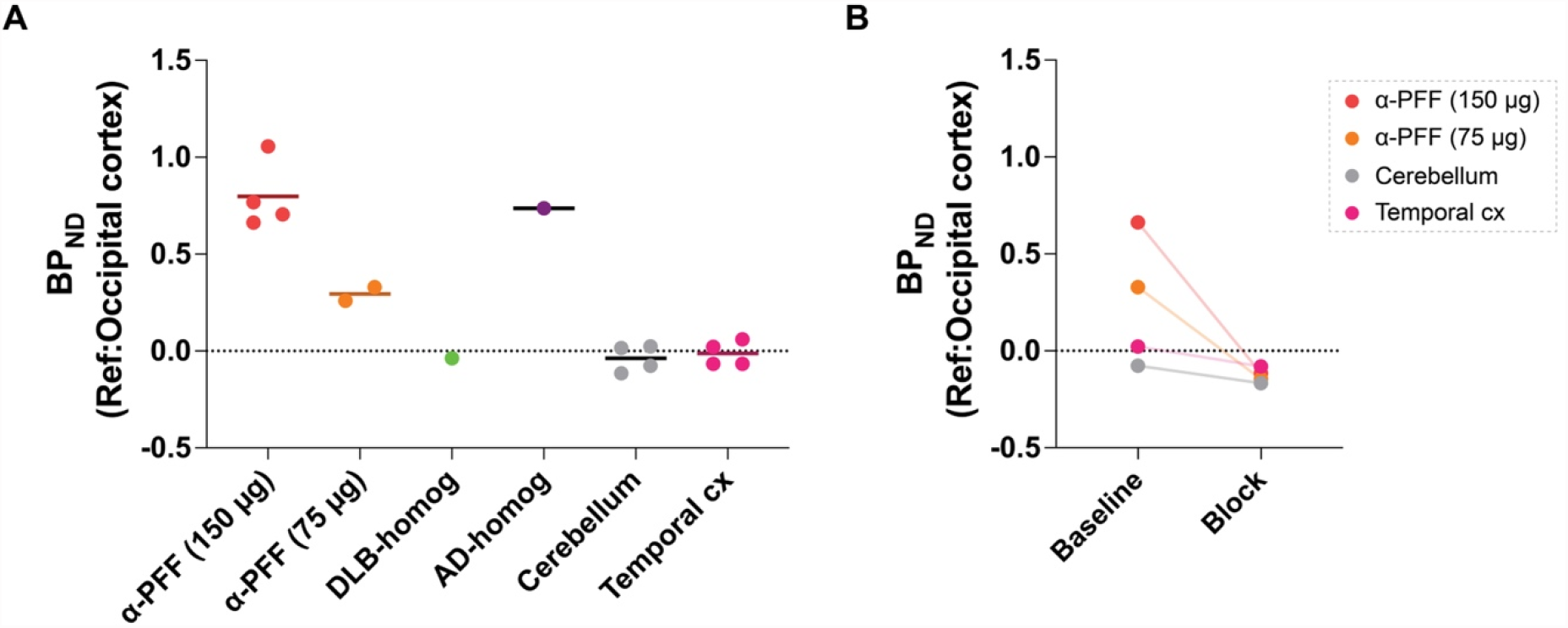
Kinetic modeling outcomes of (d_3_)-[^11^C]MODAG-001. A) BP_ND_ as determined with the non-invasive Logan graphical analysis using the occipital cortex as a reference region, in the injected brain regions, temporal cortex, and cerebellum. Retest and block are not included. B) BP_ND_ at baseline after (d_3_)-[^11^C]MODAG-001 blocking.

### Kinetic modeling of (d_3_)-[^11^C]MODAG-001

BP_ND_ in different brain regions are shown in Figure 3A. BP_ND_ in the 150 µg α-PFF regions was 0.78 ± 0.1 (mean±SD, n = 4) while in the 75 µg regions, BP_ND_ α-PFF injected regions was 0.29 (n = 2), showing a dose-dependent effect of (d_3_)-[^11^C]MODAG-001 binding to the α-PFF. BP_ND_ in the AD homogenate region was 0.73, in the same order as the 150 µg α-PFF. The DLB homogenate region, cerebellum, and temporal cortex had BP_ND_ values close to zero (Figure 3A). The (d_3_)-[^11^C]MODAG-001 test-retest scan on the same day showed a -6.2% change in BP_ND_ (Supplementary figure 2). Pretreatment with MODAG-001 resulted in a reduction in regional binding levels such that they became comparable to the reference regions. In the pig that underwent a baseline-block study, we observed >100% occupancy in the α-PFF injected regions. A modest reduction in binding was also observed in the temporal cortex and cerebellum (Figure 3B).

## Discussion

PET neuroimaging of aggregated protein has proved critical for diagnosing and monitoring disease progression and treatment evaluation in neurodegenerative diseases with amyloid-□ and tau pathology [29, 30]. The ability to detect and quantify α-synuclein aggregates in the living human brain would be a milestone achievement for the research of PD and other α-synucleinopathies [10, 31]. Due to its high affinity to α-synuclein and favorable binding in rodent models, [^11^C]MODAG-001 and its analogs are currently some of the most promising radioligands for α-synuclein neuroimaging [12, 19].

To the best of our knowledge, this is the first time (d_3_)-[^11^C]MODAG-001 has been tested in a higher species and shown promising translational results. We evaluated (d_3_)-[^11^C]MODAG-001 in a pig model of intracerebral injection of α-PFF and postmortem human AD and DLB brain homogenates. We see high brain uptake and quick-wash out of the radioligand in the brain. The pharmacokinetics in healthy mice and the α-PFF rat model were comparable to that in pigs [12]. We saw a relatively high uptake of the radioligand in the α-PFF regions at micromolar concentrations, with a dose-dependent response with 150 µg (415 µM) and 75 µg (208 µM) injections (Figure 1-3).

Kuebler et al. tested both [^11^C]MODAG-001 and the deuterium incorporated (d_3_)-[^11^C]MODAG-001 [12]; deuterium incorporation was meant to improve the pharmacokinetic and metabolic profile of the radioligand [32]. Notably, we observed faster metabolism in the pigs than what was observed in the mice, which are much smaller mammals [33]. The results showed that ∼10% parent fraction remained 15 min post-injection in the pigs, compared to ∼30% parent fraction in mice. Radio-HPLC on brain homogenate (non-perfused) from a pig euthanized at 15 min showed ∼50% parent fraction; in contrast, mice showed on average ∼90% parent fraction after 15 min (Supplementary Figure 1, Supplementary Table 1).

We performed the non-invasive kinetic modeling with the occipital cortex as a reference region since we previously have shown in our pig model that the occipital cortex has similar tissue properties as saline-injected target regions and that these are not affected by the intracerebral injection [22].

Due to the lack of other high-affinity molecules, an unlabelled MODAG-001 block scan was our best option to examine the signal specificity. Pretreatment of 1 mg/kg of MODAG-001 leads to complete blocking of the specific (d_3_)-[^11^C]MODAG-001 binding in the α-PFF injected region. We observe a very high blocking percentage with values above 100% (using BP_ND_ values from reference modeling), although these estimates are based on only one pig and likely prone to noise.

We also see high uptake in the amyloid-□ and tau-rich AD homogenates but no significant uptake in the DLB homogenate region (Figure 1 and 3); this is remarkable since DLB is considered to have a pure α-synuclein pathology. Ideally, a radioligand should have high α-synuclein selectivity for it to distinguish α-synuclein aggregates from amyloid-□ and tau aggregates [10, 34]. Several things make us less enthusiastic about the prospect of (d_3_)-[^11^C]MODAG-001 as a radioligand in human studies: (d_3_)-[^11^C]MODAG-001 did not display high binding in the DLB homogenate region; this could be due to low concentrations of aggregated α-synuclein, as is most often seen in human pathology, especially at early disease stages. This null-finding could also be due to the hypothesized difference in pathological morphology in pure α-synuclein DLB subjects [35]. (d_3_)-[^11^C]MODAG-001 was also not very selective for α-synuclein and had significant binding to the AD homogenate region. This observation is also on par with previous autoradiography studies where the highest uptake was noted in human brain sections with AD [12]. Improving the signal-to-background ratio and selectivity will be critical for the further development of the tracer, and this work is currently ongoing [12].

The intracerebral protein injection model used in the current study also comes with a set of limitations. Since the intracerebral injections are done a few hours prior to scanning, it is unlikely that protein aggregates enter into the brain cells, which does not mimic the intracellular inclusions seen in α-synucleinopathies well [3, 5]. The concentration of the α-PFF in the model is much higher than that of diseased brains, where α-synuclein is found to be at nanomolar concentration [12, 36]. This particular setup allowed us to show proof of concept for α-synuclein aggregate detecting radioligands. The signal-to-background ratio of (d_3_)-[^11^C]MODAG-001 makes it challenging to detect pathologically relevant α-synuclein, i.e., at nanomolar concentrations.

In spite of the poor specificity and relatively modest signal-to-background ratio, we believe that (d_3_)-[^11^C]MODAG-001 with its high affinity for α-synuclein is a suitable lead molecule for further radioligand development and evaluation.

## Conclusions

We demonstrate in vivo detection of α-PFF in pigs using (d_3_)-[^11^C]MODAG-001, which has previously only been shown using a similar α-PFF injection rat model. The radioligand shows excellent brain kinetics and test-retest variability. Although (d_3_)-[^11^C]MODAG-001 displays low specificity towards α-synuclein and a potential passage of radiometabolites through the blood-brain barrier, it shows promise as a lead tracer for further radiotracer development.

## List of abbreviations

α-PFF: α-synuclein preformed fibrils
AD: Alzheimer’s disease
BP_ND_: binding potential non-displaceable
DLB: dementia with Lewy bodies
HRRT: high-resolution research tomograph
IV: intra-venous
IM: intra-muscular
mPFC: medial prefrontal cortex
MSA: multiple system atrophy
PD: Parkinson’s disease
PET: positron emission tomography
R-HPLC: radio-high performance liquid chromatography
SRTM: simplified reference tissue model
SUV: standardized uptake values
TAC: time-activity curve

## Declarations

### Ethics approval

All animal procedures were performed in accordance with the European Commission’s Directive 2010/63/EU, as well as the ARRIVE guidelines, and were approved by the Danish Council of Animal Ethics (Journal no. 2017-15-0201-01375).

### Consent for publication

Not applicable

### Availability of data and material

All data, including R scripts, is available at a GitHub repository (https://github.com/nakulrrraval/Protien_inj_pig_model_MODAG001). All other requests are directed to this article’s corresponding or first author.

### Competing interests

Lundbeck A/S, Denmark provided the α-synuclein preformed fibrils as part of the European Union’s Horizon 2020 research and innovation program under the Marie Sklodowska-Curie grant agreement No. 813528. However, they had no other financial interests in the project. GMK received honoraria as a speaker and consultant for Sage Pharmaceuticals/Biogen and Sanos A/S. All other authors declare no conflict of interest.

### Funding

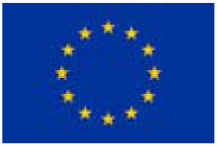

This project has received funding from the European Union’s Horizon 2020 research and innovation program under the Marie Skłodowska-Curie grant agreement No. 813528. This project also received funding from Parkinsonforeningen, Denmark (R16-A247). Pontus Plavén Sigray was supported by the Lundbeck Foundation (R303-2018-3263). Vladimir Shalgunov was supported by the Lundbeck Foundation (R303-2018-3567), the Novo Nordisk Foundation (grant agreement no. NNF18SA0034956), and BRIDGE – Translational Excellence Program at the Faculty of Health and Medical Sciences, University of Copenhagen.

### Authors’ contribution

NRR, MMH, HDH, PPS, GMK: conceptualization and design. NRR, CAM, EEB, LMJ, HDH: surgical setup and PET scanning. VS, AN, UMB: compound synthesis, radiochemistry, and HPLC analysis. NRR, VS, AN, UMB, MJ, PPS: analysis and software. NRR, MMH, HDH, GMK: resources. NR, HDH, PPS, GMK: data curation. LMJ, MMH, HDH, PPS, GMK: supervision. NRR: preparation of manuscript draft including figures. NRR, CAM, VS, AN, UMB, EEB, MJ, LMJ, MMH, HDH, PPS, GMK: manuscript review and editing. NRR, MMH, GMK: funding acquisition. All authors have read and agreed to the current version of the manuscript.

## Acknowledgments

We want to thank Lundbeck A/S, Valby, Denmark, for providing the α-synuclein preformed fibrils. This research project received human brain tissue from the Neuropathology Core of the Emory Center for Neurodegenerative Disease; we are grateful for their support. We would sincerely like to thank the staff and veterinarians at EMED, Panum, København University, and the PET and cyclotron unit at Rigshospitalet. Further, we would like to thank Ran Sing Saw, Marko Rosenholm, and Natalie Beschorner for their help during PET scans. We also extend our thanks to Ran Sing Saw for scientific discussions.

## Notes

https://github.com/nakulrrraval/Protein_inj_pig_model_MODAG001

