## Supplementary Data for "Evaluation of the α-synuclein PET radiotracer (d_3_)-[^11^C]MODAG-001 in pigs"

Supplementary information

### Supplementary Figures

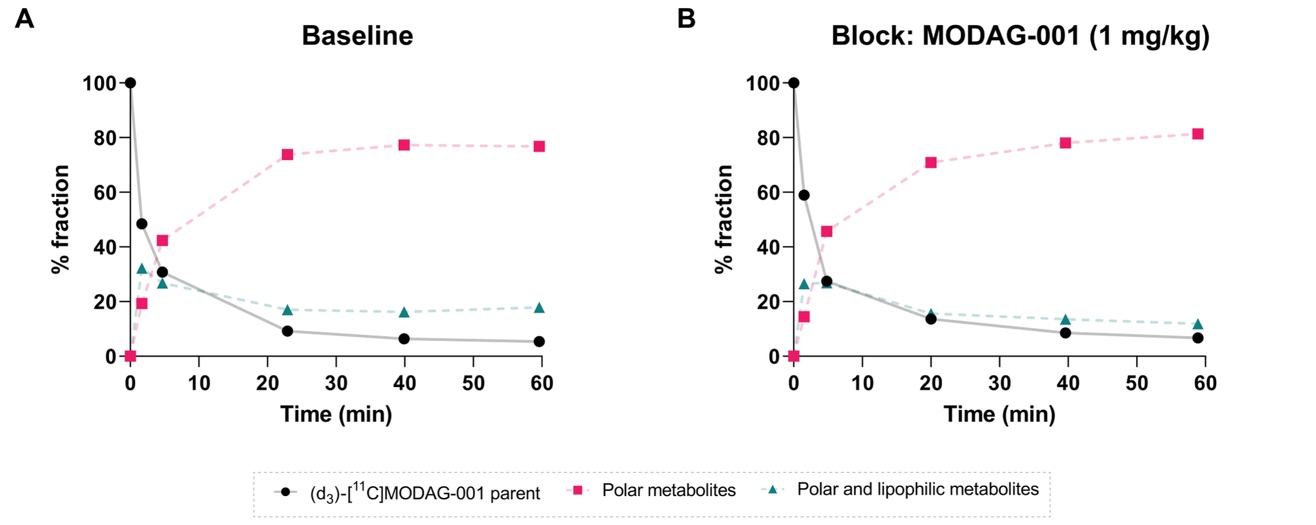

**Supplementary Figure 1. (d_3_)-[^11^C]MODAG-001 parent and metabolites.** Time course of the percentage of (d_3_)-[^11^C]MODAG-001 parent and metabolites in pig arterial plasma (n = 1 x 2 scans). The bold black line shows the course of the parent fraction over 60 min while the dashed pink and green line shows the course of the radiometabolite seen in the pigs.

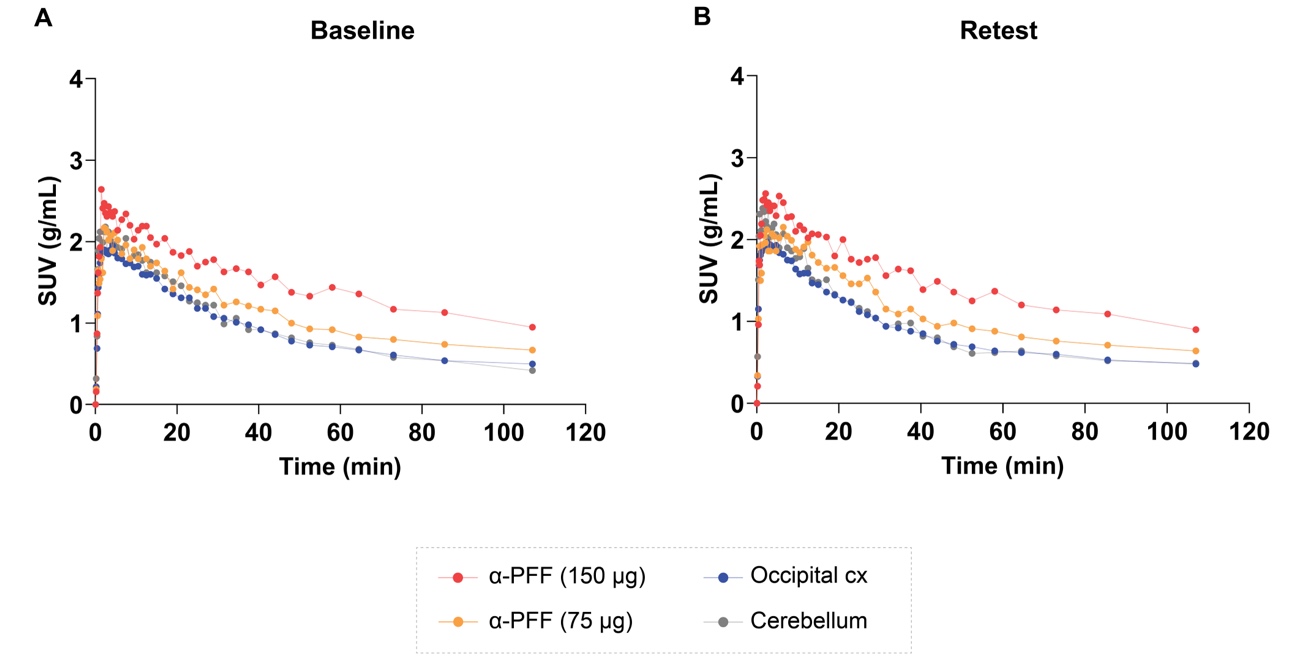

**Supplementary Figure 2. Test-retest TACs** from a pig injected with 150 µg and 75 µg α-synuclein-preformed-fibrils (α-PFF) that received two (d_3_)-[^11^C]MODAG-001 as a test (A) and retest/rescan (B).

### Supplementary Data

#### Synthesis of reagents

All reagents and solvents were dried prior to use according to standard methods. Commercial reagents were used without further purification. Analytical TLC was performed using silica gel 60 F254 (Merck) with detection by UV absorption and/or by charring following immersion in a 7% ethanolic solution of sulfuric acid or KMnO_4_-solution (1.5 g of KMnO_4_, 10 g K_2_CO_3_, and 1.25 mL 10% NaOH in 200 mL water). Purification of compounds was carried out by column chromatography on silica gel (40-60 μm, 60 Å) or employing a CombiFlash NextGen 300+ (Teledyne ISCO). ^1^H and ^13^C NMR spectra were recorded on Brucker (400 and 600 MHz instruments), using Chloroform-d, Methanol-d_4_ or DMSO-d_6_ as deuterated solvent and with the residual solvent as the internal reference. For all NMR experiments the deuterated solvent signal was used as the internal lock. Chemical shifts are reported in δ parts per million (ppm). Coupling constants (J values) are given in Hertz (Hz). Multiplicities of ^1^H NMR signals are reported as follows: s, singlet; d, doublet; dd, doublet of doublets; ddd, doublet of doublets of doublets; dt, doublet of triplets; t, triplet; q, quartet; m, multiplet; br, broad signal. NMR spectra of all compounds are reprocessed in MestReNova software (version 12.0.22023) from original FID’s files. Mass spectra analysis was performed using MS-Acquity-A: Waters Acquity UPLC

with QDa-detector.

###### 4-[3-(4-Dimethylaminophenyl)-1*H*-pyrazol-5-yl]-2-bromopyridine (MODAG-001)

To a solution of 1-[4-(dimethylamino)phenyl]ethanone (490 mg, 3.00 mmol) and methyl 2-bromopyridine-4-carboxylate (843 mg, 3.9 mmol) in DMSO (7.5 mL) and THF (1.9 mL) sodium hydride (60 % in oil, 3.9 mmol, 156 mg) was added, and the reaction mixture was stirred at 20 °C for 15 hours. The reaction mixture was poured into 50 mL of an ice water and 1 M phosphate buffer, p*H* 7 (10 mL), stirred for one hour, the resulting precipitate was filtered off, washed with water (2 ´10 mL), methanol (10 mL), hexane (10 mL), and dried on the air to obtain a crude intermediate 1-(2-bromopyridin-4-yl)-3-[4-(dimethylamino)phenyl]propane-1,3-dione as an orange solid. To a suspension of this crude intermediate in THF (20 mL) hydrazine hydrate (292 µL, 300 mg, 6 mmol) was added. The reaction mixture was stirred at 70 °C for five hours, cooled and concentrated *in vacuo*. The residue was suspended in methanol (10 mL), filtered off, washed with cold methanol (2´5 mL), recrystallized from *n*-butanol (10 mL) and *N*,*N*-dimethylformamide (0.2 mL), and dried to afford 0.47 g (46%) of (4-[3-(4-dimethylaminophenyl)-1*H*-pyrazol-5-yl]-2-bromopyridine as a light pink solid. ^1^H NMR (600 MHz, DMSO) δ 13.45 (d, J = 2.0 Hz, 1H), 8.40 (d, J = 5.1 Hz, 1H), 8.02 (d, J = 1.4 Hz, 1H), 7.86 (dd, J = 5.1, 1.4 Hz, 1H), 7.62 (d, J = 8.8 Hz, 2H), 7.25 (d, J = 2.0 Hz, 1H), 6.81 (d, J = 8.9 Hz, 2H), 2.96 (s, 6H); ^13^C NMR (101 MHz, DMSO) δ 151.27, 150.76, 147.87, 145.32, 144.78, 142.70, 126.60, 123.52, 119.55, 116.92, 112.71, 99.80, 40.37.

###### *Tert*-butyl (4-acetylphenyl)carbamate

A mixture of 4-aminoacetophenone (2.00 g, 14.79 mmol) and di-tert-butyl dicarbonate (3.52 g, 16.13 mmol) in THF (40 mL) was refluxed for 5 h. After dilution with diethyl ether (45 mL), the mixture was washed sequentially with saturated aqueous citric acid solution (20 mL), 1N NaHCO_3_ (30 mL), and brine (30 mL). The organic layer was dried over anhydrous MgSO_4_ and evaporated to give white crystals. The compound was recrystallized from EtOAc to give 3.2 g (92%) of the desired product. Rf = 0.27 (80/20 n-heptane/EtOAc); ^1^H NMR (400 MHz, CDCl_3_) δ 7.90 (d, J = 8.5 Hz, 2H), 7.45 (d, J = 8.5 Hz, 2H), 6.80 (s, 1H), 2.55 (s, 3H), 1.52 (s, 9H); ^13^C NMR (101 MHz, CDCl_3_) δ 196.90, 152.18, 142.95, 131.85, 129.84, 117.42, 81.28, 28.27, 26.35.

###### *Tert*-butyl (4-acetylphenyl)(methyl-d_3_)carbamate

A mixture of tert-butyl N-(4-acetylphenyl)carbamate (2.1 g, 8.92 mmol) in anhydrous THF (50 mL) under nitrogen was cooled to 0 °C, and sodium hydride (0.43 g, 10.71 mmol, 60 wt % dispersion in mineral oil) was added portion-wise over the course of 10 min. After 20 min, CD_3_I (1.39 mL, 22.31 mmol) was added, and the white suspension was warmed to room temperature and stirred until it formed an amber solution. After completion, the reaction was quenched with water (50 mL) and extracted with CH_2_Cl_2_ (3 x 20 mL). Purification by flash chromatography (90/10 n-heptane/EtOAc) afforded 1.71 g (76%) of the desired product as a colorless oil. Rf = 0.38 (80/20 n-heptane/EtOAc); ^1^H NMR (400 MHz, CDCl_3_) δ 7.94 (d, J = 8.5 Hz, 2H), 7.38 (d, J = 8.6 Hz, 2H), 2.60 (s, 3H), 1.50 (s, 9H); ^13^C NMR (101 MHz, CDCl_3_) δ 197.08, 154.11, 148.05, 133.43, 128.84, 124.29, 81.13, 36.31 (m), 28.29, 26.50.

###### 1-(4-((Methyl-d_3_)amino)phenyl)ethan-1-one

To a solution of the tert-butyl (4-acetylphenyl)(methyl-d_3_)carbamate (2.05 g, 8.12 mmol) in CH_2_Cl_2_ (20 mL) was added TFA (6.24 mL, 81.24 mmol). The solution was stirred at rt for 12 hours. The reaction was quenched with a saturated solution of NaHCO_3_ (20 mL) and the organic phase separated. Evaporation under reduced pressure afforded 1.22 g (99%) of the desired product as a magenta solid. Rf = 0.25 (70/30 n-heptane/EtoAc); ^1^H NMR (400 MHz, CDCl_3_) δ 7.82 (d, J = 8.8 Hz, 2H), 6.55 (d, J = 8.8 Hz, 2H), 4.10 (s, 1H), 2.49 (s, 3H); ^13^C NMR (101 MHz, CDCl_3_) δ 196.45, 153.15, 130.78, 126.56, 111.09, 29.36 (m) 25.99.

###### 1-(4-(Methyl(methyl-d_3_)amino)phenyl)ethan-1-one

To a mixture of 1-(4-((methyl-d_3_)amino)phenyl)ethan-1-one (1.00 g, 6.57 mmol) and K_2_CO_3_ (2.27 g, 16.42 mmol) in acetone (50 mL) was added CH_3_I (1.63 mL, 26.28 mmol) was added, and the white suspension was stirred for 12 hours at room temperature. After completion, the reaction was quenched with water (60 mL) and extracted with CH_2_Cl_2_ (3 x 40 mL). Purification by flash chromatography (80/20 n-heptane/EtOAc) afforded 1.05 g (96%) of the desired product as a white solid. Rf = 0.35 (70/30 n-heptane/EtOAc); ^1^H NMR (400 MHz, CDCl_3_) δ 7.88 (d, J = 9.0 Hz, 2H), 6.71 (d, J = 9.0 Hz, 2H), 3.06 (s, 3H), 2.51 (s, 3H); ^13^C NMR (151 MHz, CDCl_3_) δ 196.40, 153.08, 130.54, 126.05, 111.13, 40.27, 39.57 (m), 26.02.

###### 4-(5-(2-Bromopyridin-4-yl)-1H-pyrazol-3-yl)-N-methyl-N-(methyl-d_3_)aniline (d3-MODAG-001 reference)

To a solution of 1-(4-(Methyl(methyl-d_3_)amino)phenyl)ethan-1-one (0.63 g, 3.79 mmol) and methyl 2-bromopyridine-4-carboxylate (1.06 g, 4.92 mmol) in DMSO (15 mL) and THF (4 mL) sodium hydride (0.19 g, 7.58 mmol, 60 wt % dispersion in mineral oil) was added, and the reaction mixture was stirred at 20 °C for 15 hours. The reaction mixture was poured into 50 mL of an ice water and 1 M phosphate buffer, pH 7 (10 mL). The resulting precipitate was filtered off, washed with water (2 x 10 mL), methanol (10 mL), hexane (10 mL), and dried on to obtain the crude dione intermediate as an orange solid. To a suspension of this crude intermediate in THF (20 mL) hydrazine hydrate (0.37 mL, 7.58 mmol) was added. The reaction mixture was stirred at 70 °C for five hours, cooled and concentrated under reduced pressure. The residue was suspended in methanol (10 mL), filtered off, washed with cold methanol and recrystallized from n-butanol (10 mL) and N,N-dimethylformamide (0.2 mL) to give 0.56 g (43%) of the desired product. ^1^H NMR (600 MHz, DMSO) δ 13.46 (s, 1H), 8.40 (d, J = 5.1 Hz, 1H), 8.03 (s, 1H), 7.86 (d, J = 5.2 Hz, 1H), 7.63 (d, J = 8.3 Hz, 2H), 7.25 (s, 1H), 6.79 (d, J = 8.3 Hz, 2H), 2.94 (s, 3H).

###### Tert-butyl (4-(3-(2-bromopyridin-4-yl)-3-oxopropanoyl)phenyl)(methyl-d_3_)carbamate

To a solution of tert-butyl N-(4-acetylphenyl)-N-(^2^H_3_)methylcarbamate (1.7 g, 6.73 mmol) and methyl 2-bromopyridine-4-carboxylate (1.89 g, 8.75 mmol) in DMSO (15 mL) and THF (4 mL) sodium hydride (0.35 g, 8.75 mmol, 60 wt % dispersion in mineral oil) was added, and the reaction mixture was stirred at 20 °C for 15 hours. The reaction mixture was poured into 50 mL of an ice water and 1 M phosphate buffer, pH 7 (10 mL). Extraction with CH_2_Cl_2_ (3 x 40 mL) afforded 3.1 g of crude. Purification by flash chromatography (90/10 n-heptane/EtOAc) afforded 1.05 g (36%) of the desired product as a yellow oil. Rf = 0.17 (80/20 n-heptane/EtOAc); ^1^H NMR (400 MHz, CDCl_3_) δ 8.53 (dd, J = 5.1, 0.7 Hz, 1H), 8.02 – 7.92 (m, 3H), 7.73 (dd, J = 5.1, 1.5 Hz, 1H), 7.44 (d, J = 8.9 Hz, 2H), 6.80 (s, 1H), 1.50 (s, 9H); ^13^C NMR (101 MHz, CDCl_3_) δ 187.86, 179.09, 154.02, 150.94, 148.47, 145.14, 143.16, 130.87, 128.05, 125.13, 124.41, 119.49, 94.07, 81.37, 35.76 (m), 28.31.

###### Tert-butyl (4-(5-(2-bromopyridin-4-yl)-1H-pyrazol-3-yl)phenyl)(methyl-d_3_)carbamate

To a suspension of tert-butyl (4-(3-(2-bromopyridin-4-yl)-3-oxopropanoyl)phenyl)(methyl-d_3_)carbamate (0.70 g, 1.60 mmol) in THF (20 mL) hydrazine hydrate (0.16 mL, 3.20 mmol) was added. The reaction mixture was stirred at 60 °C for five hours, cooled and concentrated under reduced pressure. The compound was purified by flash chromatography (98/2 CH_2_Cl_2_/MeOH) to yield 0.68 g (98%) of the desired compound as a white foam. Rf = 0.35 (95/5 CH_2_Cl_2_/MeOH); ^1^H NMR (400 MHz, CDCl_3_) δ 10.88 (s, 1H), 8.40 (d, J = 5.2 Hz, 1H), 7.92 (s, 1H), 7.67 (dd, J = 5.2, 1.5 Hz, 1H), 7.55 (d, J = 8.5 Hz, 2H), 7.35 (d, J = 8.5 Hz, 2H), 6.87 (s, 1H), 1.50 (s, 9H); ^13^C NMR (101 MHz, CDCl_3_) δ 154.64, 150.50, 144.41, 142.93, 125.83, 125.73, 124.15, 119.11, 100.97, 80.97, 53.42, 36.23 (m), 28.37.

###### 4-(5-(2-Bromopyridin-4-yl)-1H-pyrazol-3-yl)-N-(methyl-d_3_)aniline (desmethyl precursor for reductive amination)

To a solution of tert-butyl (4-(5-(2-bromopyridin-4-yl)-1H-pyrazol-3-yl)phenyl)(methyl-d_3_)carbamate (0.60 g, 1.39 mmol) in CH_2_Cl_2_ (5 mL) was added TFA (1.06 mL, 13.90 mmol). The reaction was stirred for 2 hours and concentrated under reduced pressure. Purification by flash chromatography (98/2 CH_2_Cl_2_/MeOH) afforded 0.42 (91%) g of the desired compound as a beige solid. Rf = 0.30 (95/5 CH_2_Cl_2_/MeOH); ^1^H NMR (400 MHz, DMSO) δ 13.29 (s, 1H), 8.31 (d, J = 5.2 Hz, 1H), 7.92 (s, 1H), 7.76 (d, J = 5.2 Hz, 1H), 7.45 (d, J = 8.2 Hz, 2H), 7.11 (s, 1H), 6.53 (d, J = 8.4 Hz, 2H), 5.84 (s, 1H).

#### Radiochemistry

###### General information

Radiochemistry was performed at the Department of Clinical Physiology, Nuclear Medicine & PET, Rigshospitalet, Denmark. [^11^C]CH_4_ was produced via the ^14^N(p,α)^11^C reaction in a gaseous target of N_2_ (+10% H_2_) with a 16 MeV proton beam in a Scanditronix MC32 cyclotron (Scandtronix Magnet AB, Vislanda, Sweden) and converted into [^11^C]CH_3_I via gas-phase iodination as described in Larsen et. al. [[1]](https://paperpile.com/c/qWybvv/ZVYZ). The time at which [^11^C]CH_4_ was delivered to the synthesis module is defined as the start-of-synthesis, while the end-of-synthesis is defined as the time at which the activity of the formulated (d_3_)-[^11^C]MODAG-001 was measured. Automated synthesis was performed on a Scansys Laboratorieteknik (Scansys Laboratorieteknik ApS, Værløse, Denmark) synthesis module housed in a hot cell. Analytical HPLC was performed on a Dionex system connected to a P680A pump, a UVD 170U UV/Vis detector, and a Scansys radiodetector. The system was controlled by Chromeleon software. Semi-preparative HPLC was performed on the built-in HPLC system in the synthesis module.

###### Synthesis of (d_3_)-[^11^C]MODAG-001

(d_3_)-[^11^C]MODAG-001 was prepared as described in [[2]](https://paperpile.com/c/qWybvv/2WoP) with minor modifications. Namely, [^11^C]CH_3_I was bubbled in a helium stream through a solution of trimethylamine N-oxide (5 mg) and 4-(5-(2-bromopyridin-4-yl)-1H-pyrazol-3-yl)-N-(methyl-d3)aniline (1 mg) in diethyl formamide (350 µL) at -20 °C. The mixture was heated to 60 °C for 5 min and subsequently cooled to 40 °C. Sodium cyanoborohydride (7-8 mg) in a mixture of diethyl formamide (60 µL) and sodium citrate buffer (100 mM pH 4.6, 0.55 mL) was added and heated to 100 °C for 5 min. After labeling, the reaction was diluted with water (2.7 mL) and purified with semipreparative HPLC (Supplementary Figure 3) on a Luna C18 column (5 µm, 100 Å, 250 mm x 10 mm, Phenomenex, Germany) with 33 % MeCN and 0.1% TFA in water at a flow rate of 4 mL/min. The HPLC fraction containing the product was diluted with 70 mL of water and loaded onto a Sep Pak C18 Plus Light cartridge (Waters, USA). The product was eluted with 0.7 mL of ethanol and diluted with 10 mL of phosphate buffer (100 mM, pH 7.2).

Radiochemical conversion (RCC) of (d_3_)-[^11^C]MODAG-001 was determined by analyzing an aliquot labeling of the reaction mixture by analytical radio-HPLC. The identity of (d_3_)-[^11^C]MODAG-001 was confirmed by co-elution with an unlabelled ^12^C-reference compound on radio-HPLC. The molar activity (Am) was determined by integrating the area of the UV absorbance peak corresponding to the radiolabeled product on the analytical HPLC chromatogram (average of 2 runs). This area was converted into mass concentration by comparison with a calibration curve for a known range of concentrations of (d_3_)-MODAG-001 reference.

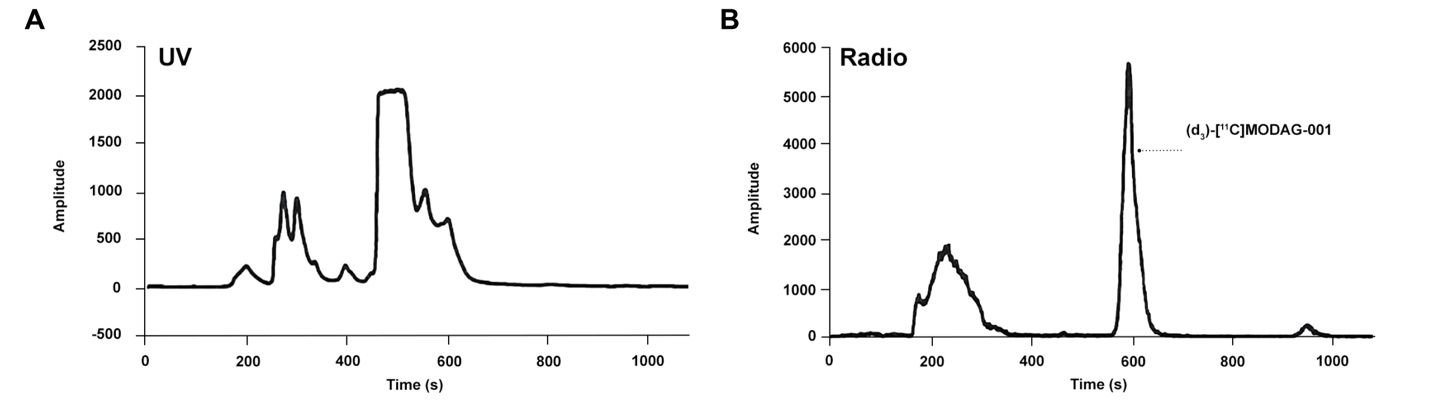

**Supplementary Figure 3. Semipreparative HPLC.** Representative UV and radio trace for (d_3_)-[^11^C]MODAG-001.

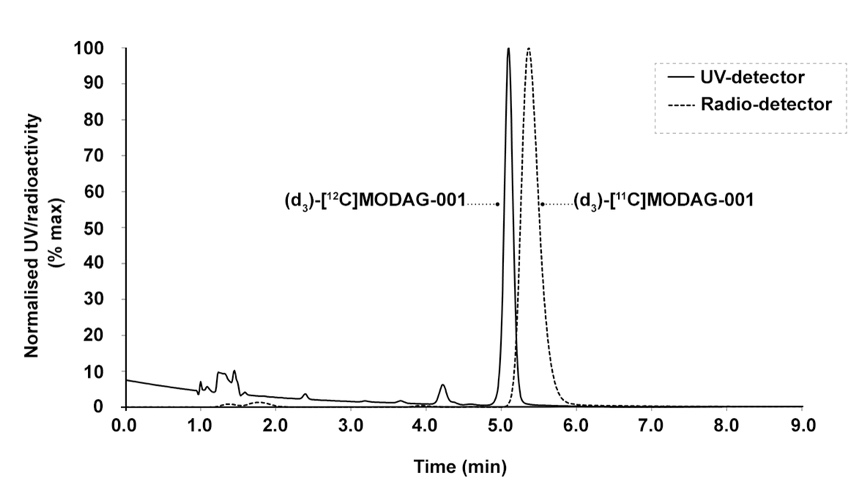

**Supplementary Figure 4. Analytical HPLC.** Representative UV and radio trace for (d_3_)-[^11^C]MODAG-001 spiked with unlabelled (d_3_)-[^12^C]MODAG-001. HPLC conditions: Luna 5μm PFP(2) 150x4.6 mm eluted with ACN/water/TFA (30/70/0.1 v/v) at 1.5 mL/min.

#### Radio-HPLC of blood and brain samples:

The HPLC system was equipped with a Luna C18(2) column (5 µm, 250 x 4.6 mm; Phenomenex, Torrance, CA, USA) eluting with 65% of 0.1% (v/v) trifluoroacetic acid in water and 35 % acetonitrile at a flow rate of 1.5 mL/min. Blood was centrifuged for 5 min at 3234 x g, 4°C. Brain tissue was weighed, homogenized in 4 volumes of PBS, and centrifuged for 5 min at 3234 x g and 4°C. Plasma and brain homogenate was precipitated with ice-cold acetonitrile (1:1) and centrifuged for 5 min at 6000 x g, 4°C. The supernatant of both samples was subsequently filtered through a syringe filter (Whatman GD/X 13 mm, PVDF membrane, 0.45 mm pore size; Frisenette ApS, Knebel, Denmark) before injection of 3 mL into the HPLC. The eluent from the HPLC system was passed through the radiochemical detector (Posi-RAM Model 4; LabLogic, Sheffield, UK) for online detection of radioactive metabolites and parent tracer. Eluents from the HPLC were collected with a fraction collector (Foxy Jr FC144; Teledyne, Thousand Oaks, CA, USA), and fractions were counted offline in a gamma well counter (2480 Wizard2 Automatic Gamma Counter, PerkinElmer, Finland). The parent fraction was determined as the percentage of the radioactivity of the parent to the total radioactivity collected.

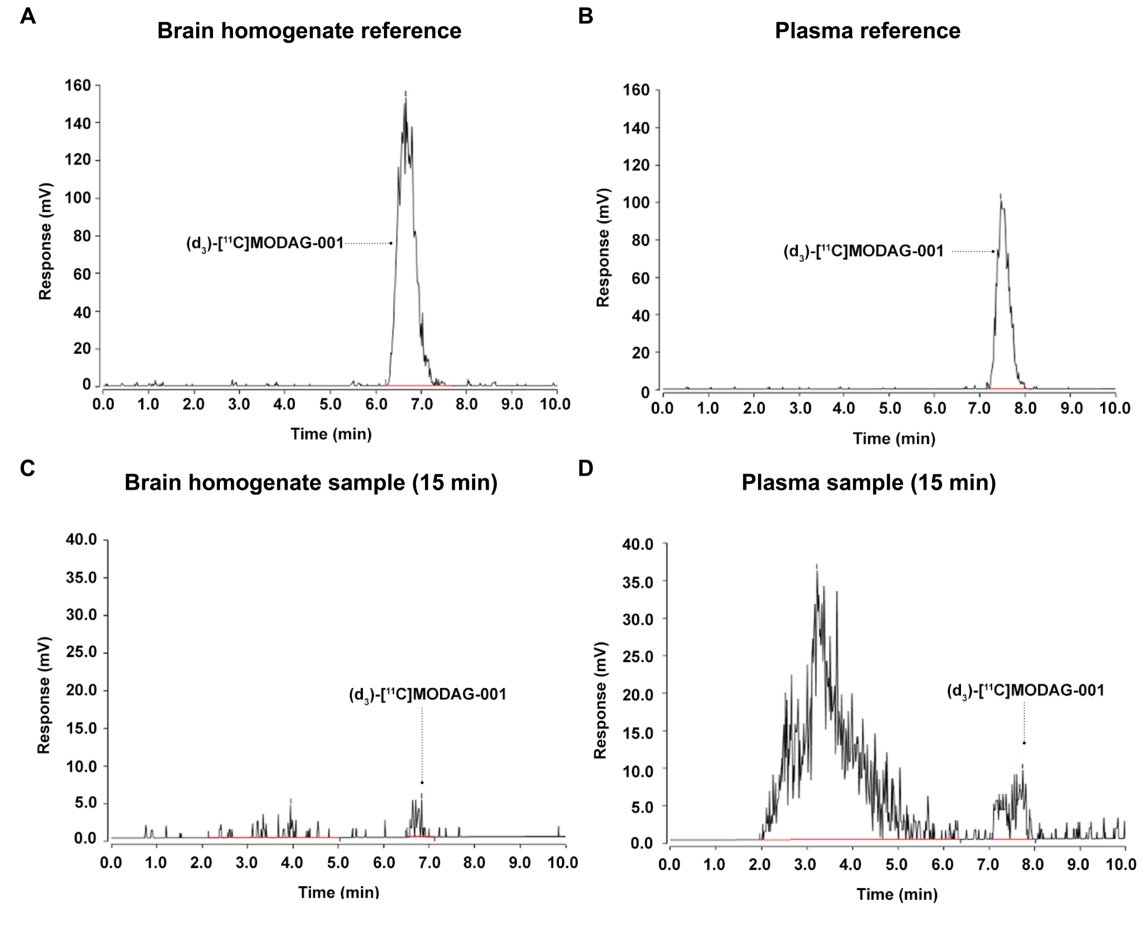

**Supplementary Figure 5. HPLC chromatograms.** Representative reference radio-chromatograms from (d_3_)-[^11^C]MODAG-001 spiked brain homogenate (A) and plasma (B). Radio-chromatograms from brain homogenate (C) and plasma (D) samples 15 min post tracer injection at euthanasia (n=1).

**Supplementary Table 1: Percentage of (d_3_)-[^11^C]MODAG-001 parent and radiometabolites.** Percentage of parent and radiometabolite calculated with radio-HPLC + gamma counting in plasma and brain at approximately 15 min post tracer injection (n = 1).

| **Sample** | **Polar Metabolites**  **(%)** | **Polar & Lipophilic Metabolites**  **(%)** | **Parent tracer**  **(%)** |
| --- | --- | --- | --- |
| Plasma *@Euthanasia*  (15 min) | 80.8 | 8.4 | 10.8 |
| Brain *@Euthanasia*  (22 min) | 30.4 | 13.6 | 56.1 |

#### Fluorescence immunostaining

Fluorescence immunostaining was performed on sections containing the injection site to validate the injection site and confirm injection of injectate. The sections were processed for standard immunohistochemistry (IHC) with α-synuclein (GTX21904-Mouse [4B12], GeneTex, Hsinchu City, Taiwan) and amyloid-ꞵ (ab252816-Rat [2E9], Abcam, Cambridge, UK) primary antibodies detected with goat anti-mouse or goat anti-rat IgG H&L (Alexa Fluor® 647) (ab150115/ab150167, Abcam, Cambridge, UK) secondary antibodies. α-synuclein containing sections were α-PFF, and dementia with Lewy bodies (DLB) homogenate injected pig brain sections while amyloid-ꞵ containing sections were Alzheimer's disease (AD) homogenate injected regions. The frozen sections were first fixed in 4% formaldehyde for 20 min. After a 10 min wash in phosphate-buffered saline (PBS), antigen retrieval was performed in a microwave oven in 25 mM sodium citrate buffer pH 7.5 for α-synuclein containing section or Tris/EDTA buffer pH 9.0 for amyloid-ꞵ containing section. Buffer temperature was raised to boiling for 5 secs, sections left in the microwave for 10 min, then 20 min under the hood. After, they were washed with PBS-TritonX100 0.4% and then incubated for 60 min with PBS + 0.4% TritonX100 and 5% bovine serum albumin (BSA). Buffer was poured off, sectioned for incubated overnight in primary antibody (α-synuclein: 0.2 ng/ml) (amyloid-ꞵ: 0.5 µg/ml) in PBS + 0.1% Tween20 at 4 °C. The next day, sectioned were washed thrice in PBS and then incubated in secondary antibody (diluted 1:200) in PBS + 0.1% Tween20 for 1 hour at room temperature. Finally, the sections were washed thrice in PBS followed by one wash in deionized H_2_O (dH_2_O). After the staining protocol, the sections were mounted in EverBrite™ Hardset Mounting Medium (Biotium, Inc., Fremont, CA, USA). Sections were imaged using an EC Plan-Neofluoar 5x/0.16 objective on an Axio Observer 7 fitted with a motorized stage and Axiocam 506mono CCD camera (Carl Zeiss, Birkerød, Denmark) to create stitched images covering large regions of interest. For Alexa Fluor® 647, an excitation of 640/30 nm, beamsplitter of 660, and emission of 690/50 filter set was used (Filter Set 50, Carl Zeiss, Birkerød, Denmark).

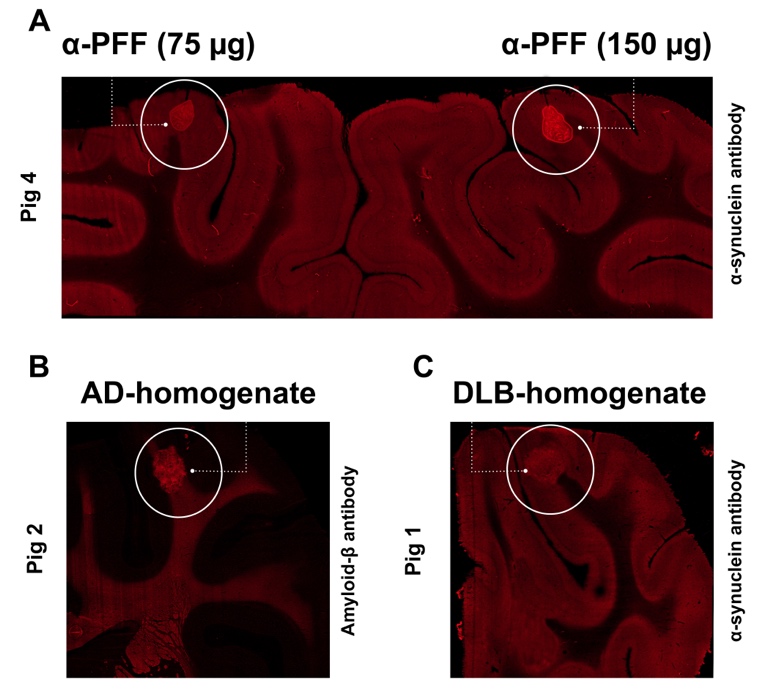

**Supplementary Figure 6. Fluorescence immunostaining.** Successful injections were ensured by immunostaining injection site containing sections. Representative examples of pig injected with A) α-PFF (right side = 150 µg, left side = 75 µg) B) AD-homogenate, and C) DLB-homogenate.
